## Supplementary material for "Genomics of Preaxostyla Flagellates Illuminates the Path Towards the Loss of Mitochondria": S4File

S4 File

A) DLP

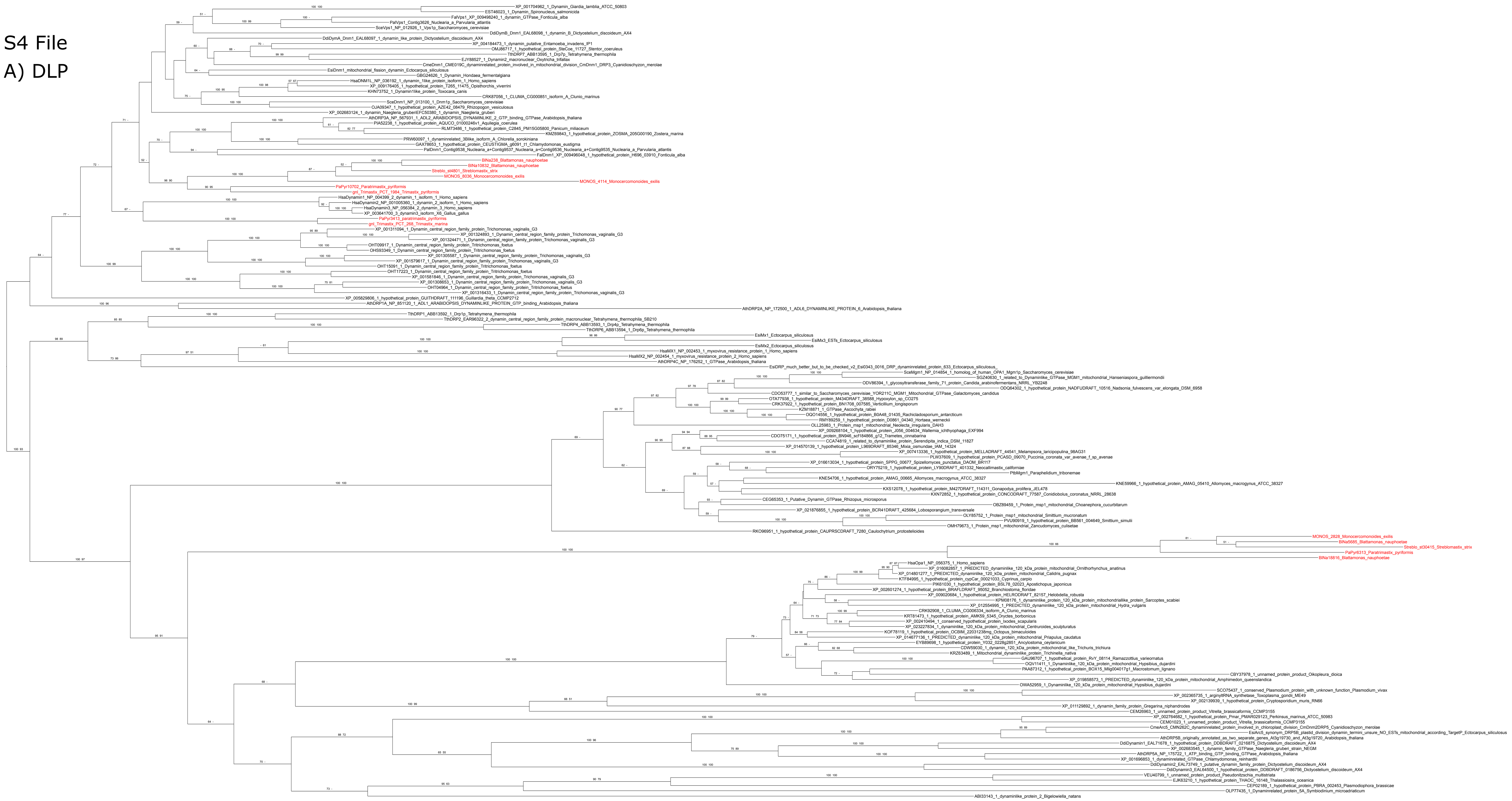

B) ME

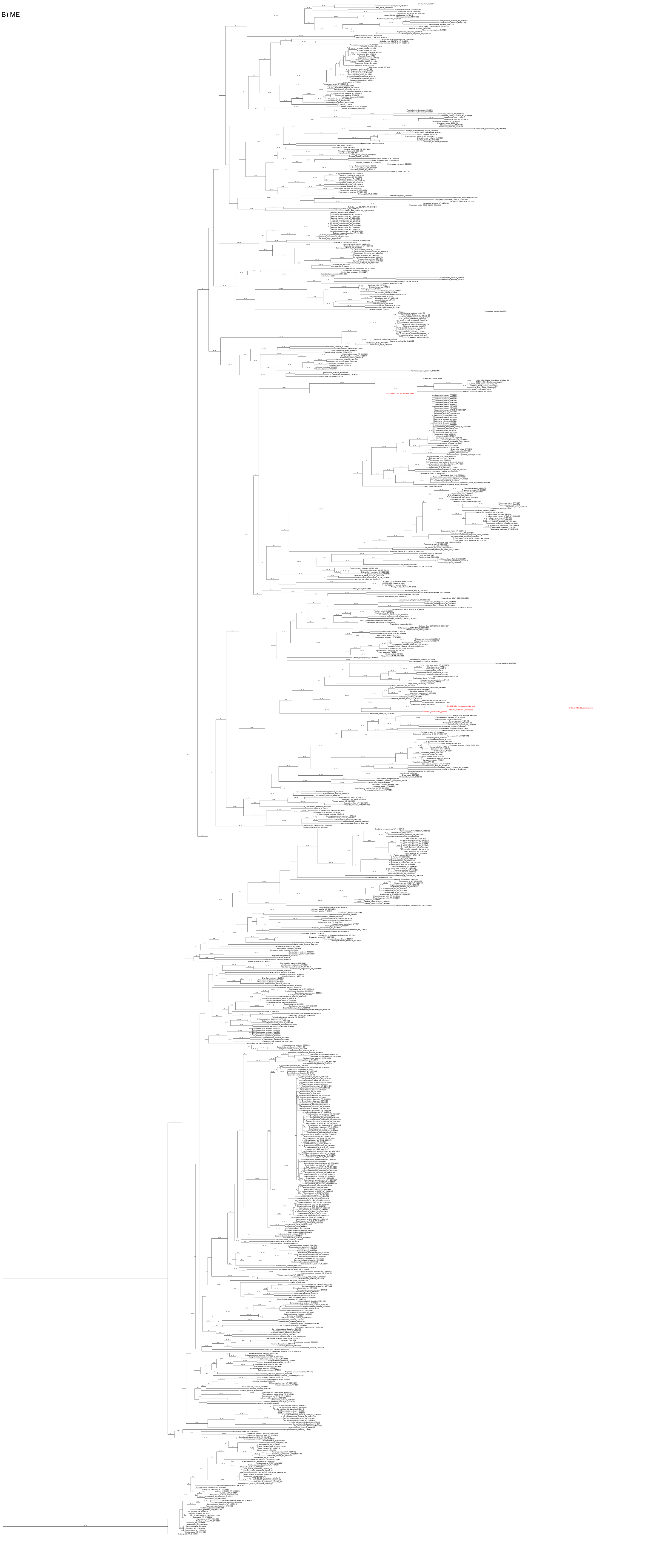

### C) PFO

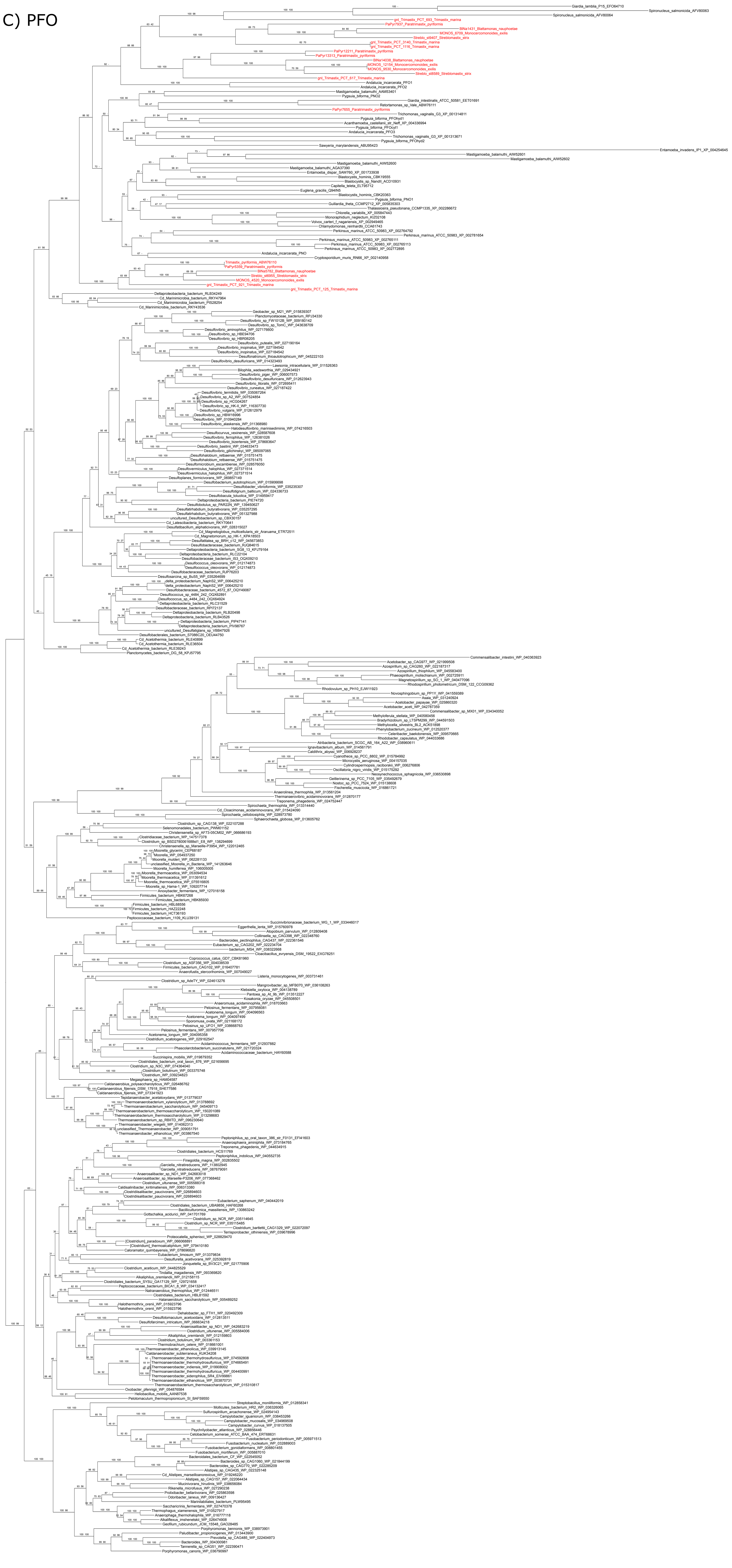

D) HydA

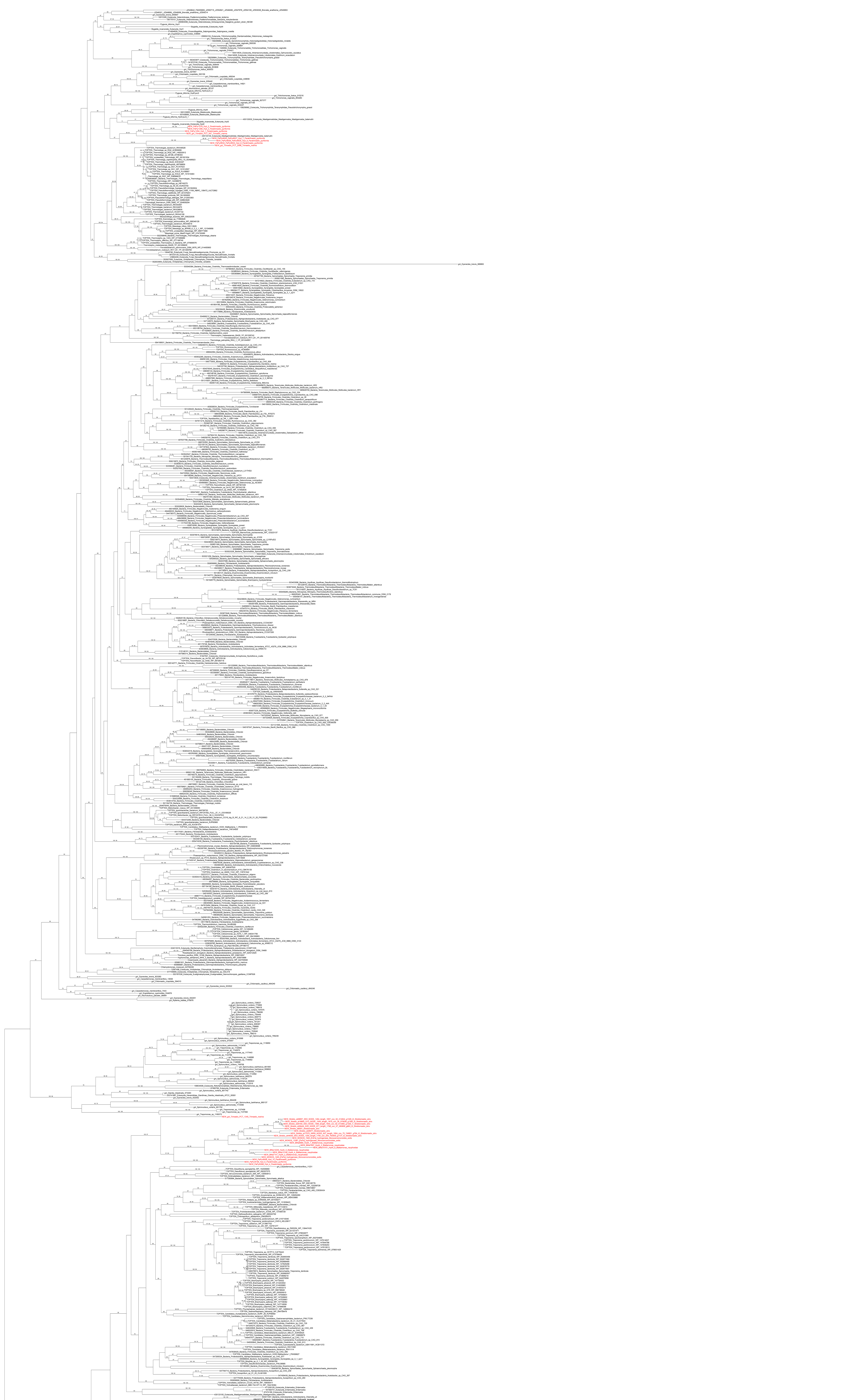

E) ACS

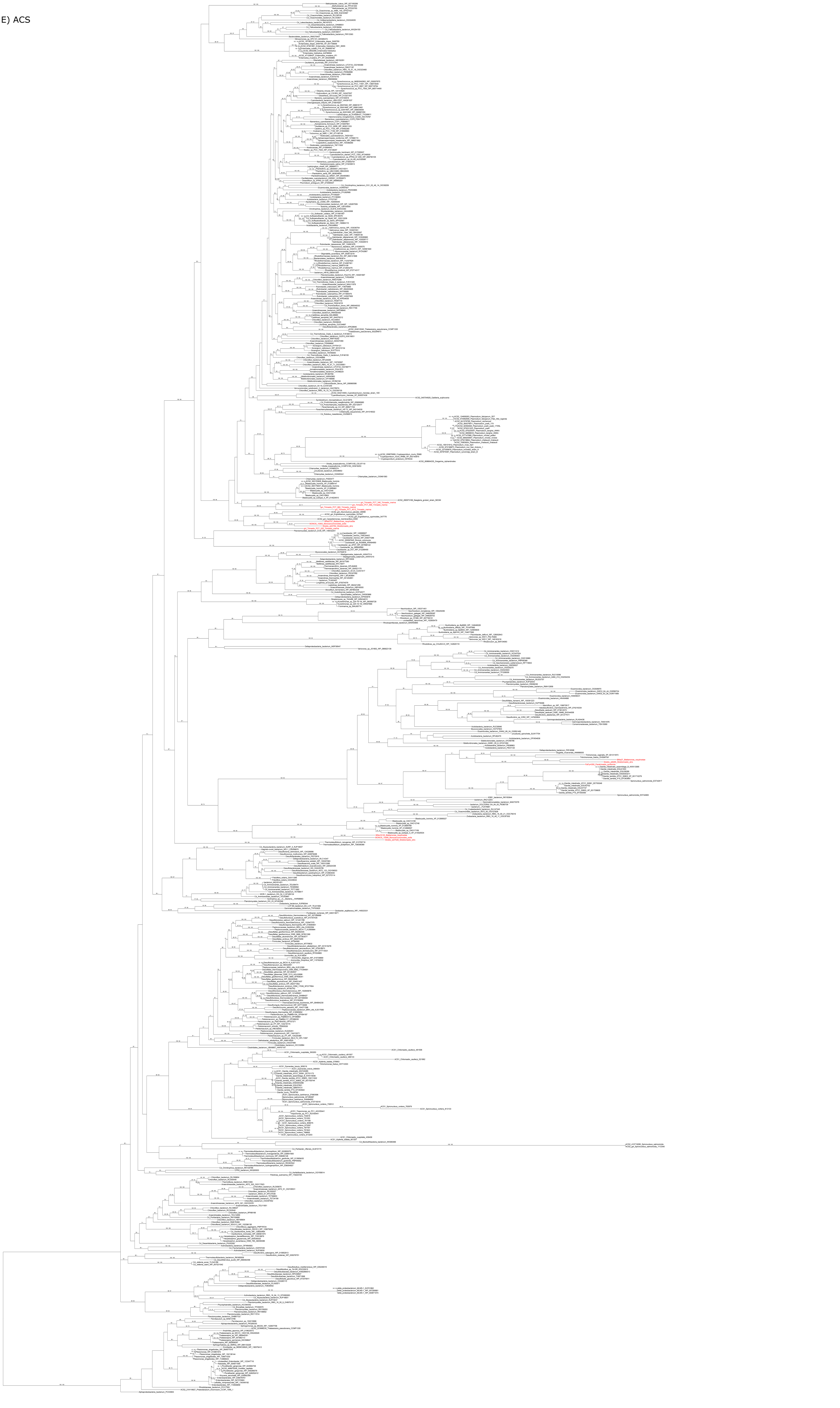

F) GCS L

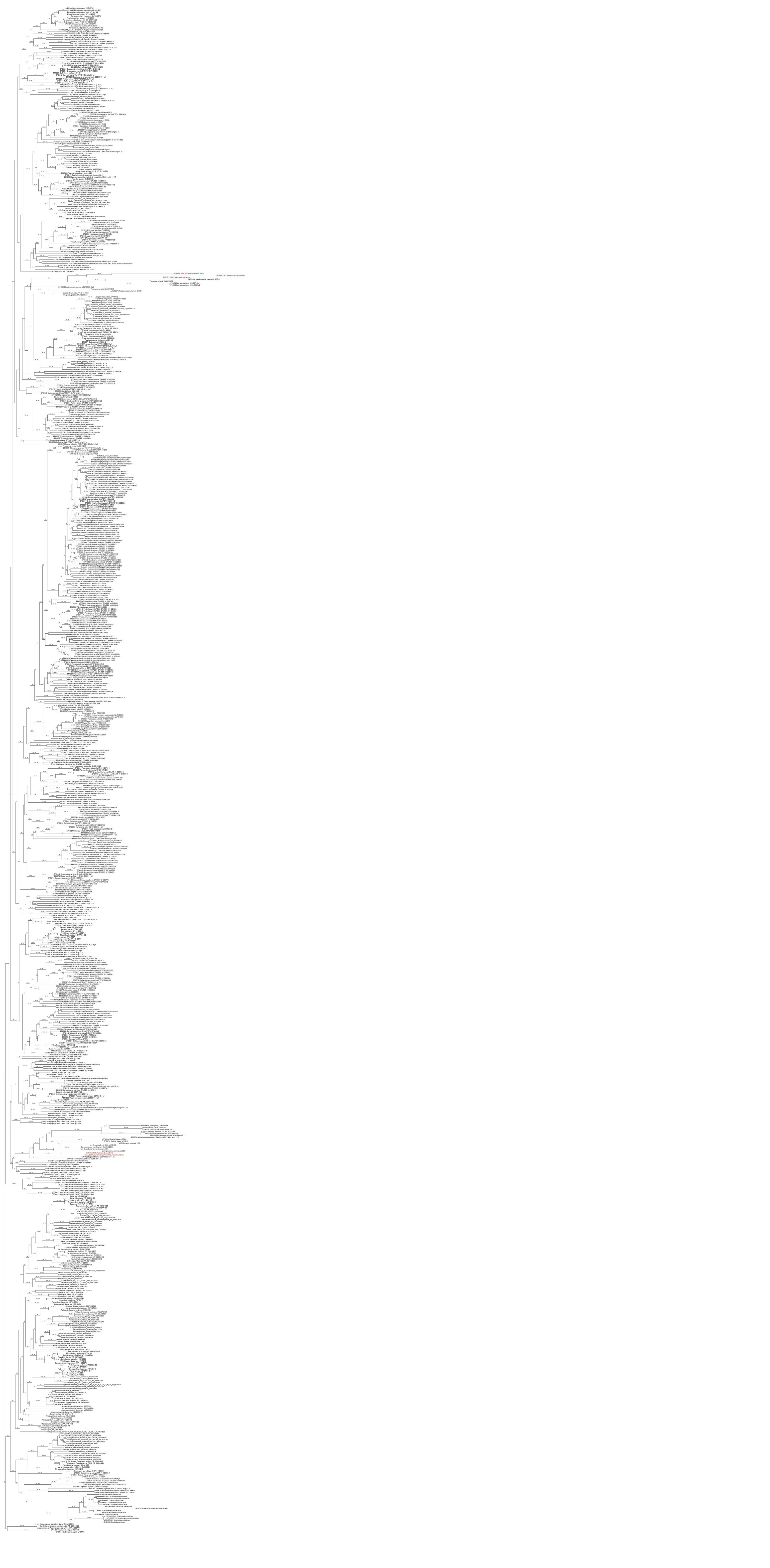

G) AAAP

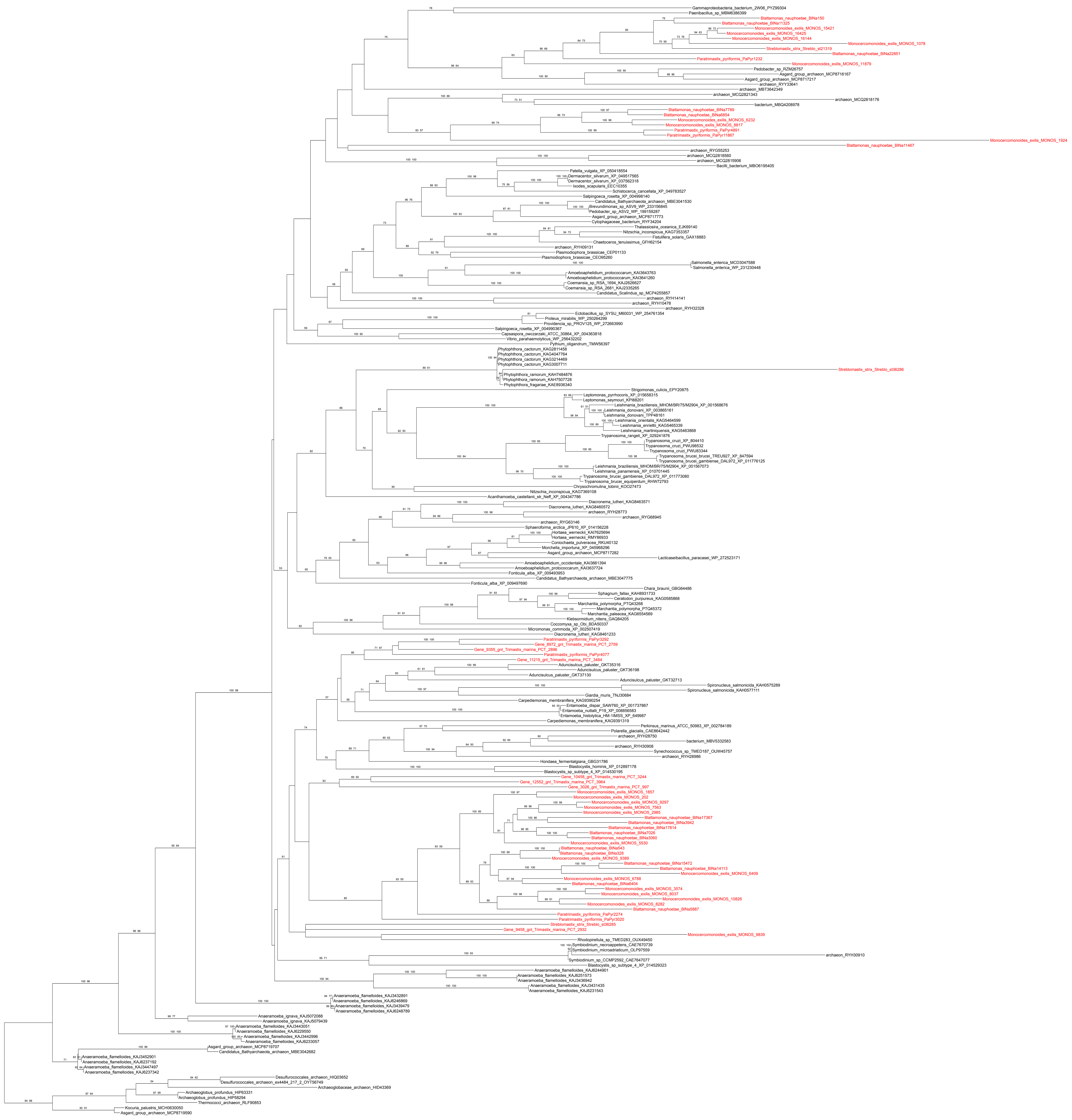

H) CTL

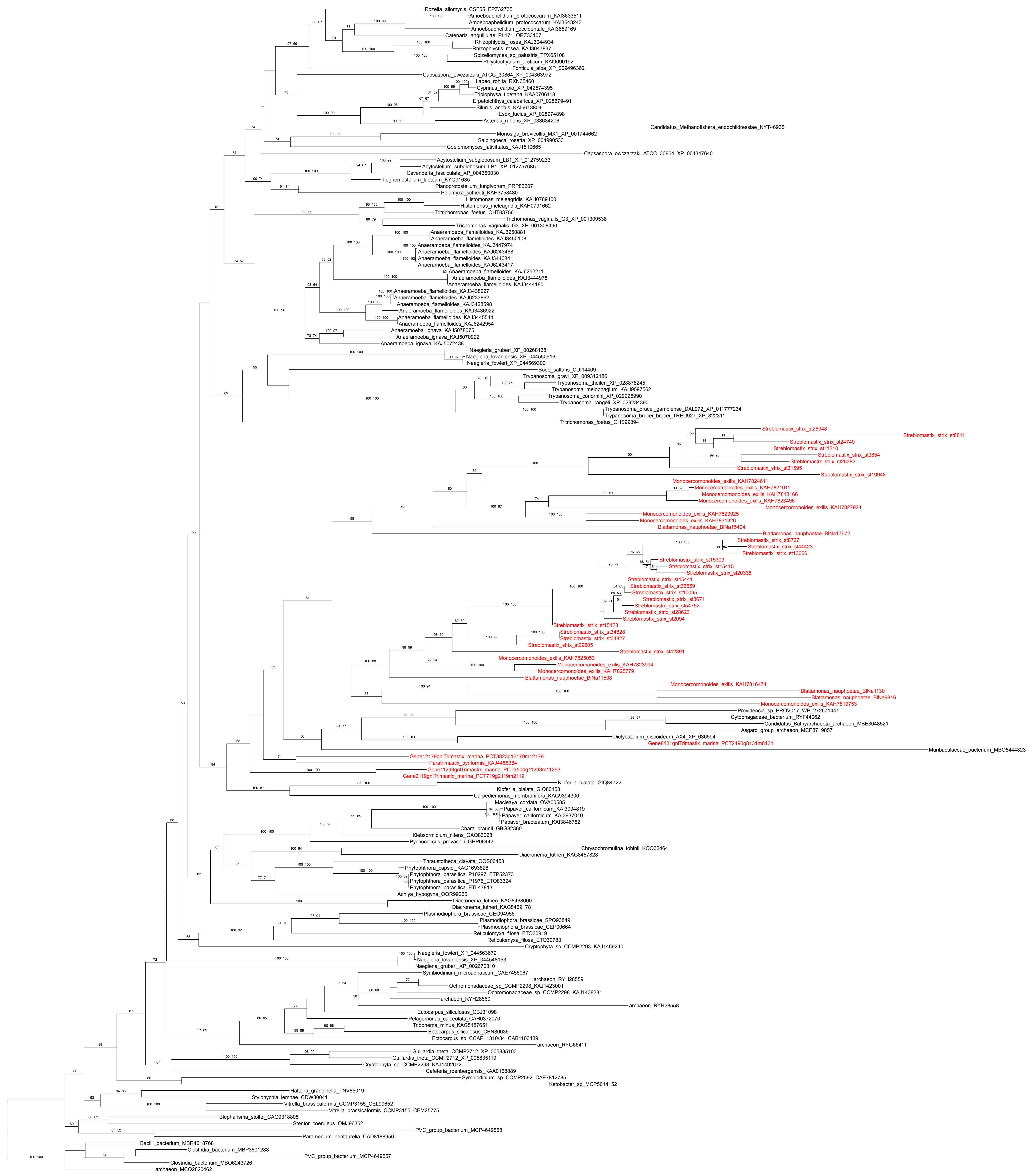

I) ENT

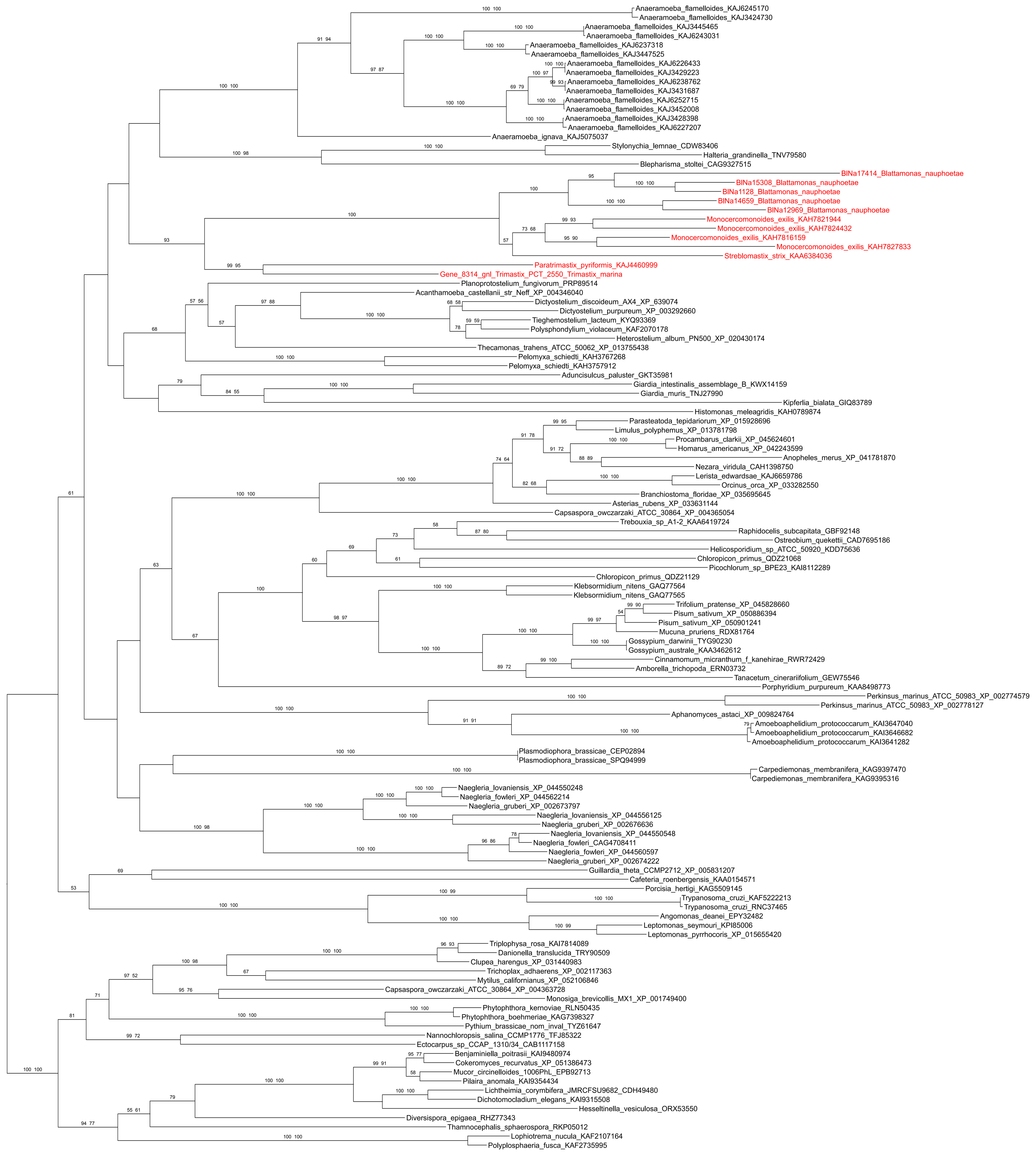

Anaeramoeba_flamelloides_KAJ6245170
Anaeramoeba_flamelloides_KAJ3424730
Anaeramoeba_flamelloides_KAJ3445465
Anaeramoeba_flamelloides_KAJ6243031
Anaeramoeba_flamelloides_KAJ6237318
Anaeramoeba_flamelloides_KAJ3447525
Anaeramoeba_flamelloides_KAJ6226433
Anaeramoeba_flamelloides_KAJ3429223
Anaeramoeba_flamelloides_KAJ6238762
Anaeramoeba_flamelloides_KAJ3431687
Anaeramoeba_flamelloides_KAJ6252715
Anaeramoeba_flamelloides_KAJ3452008
Anaeramoeba_flamelloides_KAJ3428398
Anaeramoeba_flamelloides_KAJ6227207
Anaeramoeba_ignava_KAJ5075037
Stylonychia_lemnae_CDW83406
Halteria_grandinella_TNV79580
Blepharisma_stoltei_CAG9327515
BlNa17414_Blattamonas_nauphoetae
BlNa15308_Blattamonas_nauphoetae
BlNa1128_Blattamonas_nauphoetae
BlNa14659_Blattamonas_nauphoetae
BlNa12969_Blattamonas_nauphoetae
Monocercomonoides_exilis_KAH7821944
Monocercomonoides_exilis_KAH7824432
Monocercomonoides_exilis_KAH7816159
Monocercomonoides_exilis_KAH7827833
Streblomastix_strix_KAA6384036
Paratrimastix_pyriformis_KAJ4460999
Gene_8314_gnl_Trimastix_PCT_2550_Trimastix_marina
Planoprotostelium_fungivorum_PRP89514
Acanthamoeba_castellanii_str_Neff_XP_004346040
Dictyostelium_discoideum_AX4_XP_639074
Dictyostelium_purpureum_XP_003292660
Tieghemostelium_lacteum_KYQ93369
Polysphondylium_violaceum_KAF2070178
Heterostelium_album_PN500_XP_020430174
Thecamonas_trahens_ATCC_50062_XP_013755438
Pelomyxa_schiedti_KAH3767268
Pelomyxa_schiedti_KAH3757912
Aduncisulcus_paluster_GKT35981
Giardia_intestinalis_assemblage_B_KWX14159
Giardia_muris_TNJ27990
Kipferlia_bialata_GIQ83789
Histomonas_meleagridis_KAH0789874
Parasteatoda_tepidariorum_XP_015928696
Limulus_polyphemus_XP_013781798
Procambarus_clarkii_XP_045624601
Homarus_americanus_XP_042243599
Anopheles_merus_XP_041781870
Nezara_viridula_CAH1398750
Lerista_edwardsae_KAJ6659786
Orcinus_orca_XP_033282550
Branchiostoma_floridae_XP_035695645
Asterias_rubens_XP_033631144
Capsaspora_owczarzaki_ATCC_30864_XP_004365054
Trebouxia_sp_A1-2_KAA6419724
Raphidocelis_subcapitata_GBF92148
Ostreobium_quekettii_CAD7695186
Helicosporidium_sp_ATCC_50920_KDD75636
Chloropicon_primus_QDZ21068
Picochlorum_sp_BPE23_KAI8112289
Chloropicon_primus_QDZ21129
Klebsormidium_nitens_GAQ77564
Klebsormidium_nitens_GAQ77565
Trifolium_pratense_XP_045828660
Pisum_sativum_XP_050886394
Pisum_sativum_XP_050901241
Mucuna_pruriens_RDX81764
Gossypium_darwinii_TYG90230
Gossypium_australe_KAA3462612
Cinnamomum_micranthum_f_kanehirae_RWR72429
Amborella_trichopoda_ERN03732
Tanacetum_cinerariifolium_GEW75546
Porphyridium_purpureum_KAA8498773
Perkinsus_marinus_ATCC_50983_XP_002774579
Perkinsus_marinus_ATCC_50983_XP_002778127
Aphanomyces_astaci_XP_009824764
Amoeboaphelidium_protococcarum_KAI3647040
Amoeboaphelidium_protococcarum_KAI3646682
Amoeboaphelidium_protococcarum_KAI3641282
Plasmodiophora_brassicae_CEP02894
Plasmodiophora_brassicae_SPQ94999
Carpediemonas_membranifera_KAG9397470
Carpediemonas_membranifera_KAG9395316
Naegleria_lovaniensis_XP_044550248
Naegleria_fowleri_XP_044562214
Naegleria_gruberi_XP_002673797
Naegleria_lovaniensis_XP_044556125
Naegleria_gruberi_XP_002676636
Naegleria_lovaniensis_XP_044550548
Naegleria_fowleri_CAG4708411
Naegleria_fowleri_XP_044560597
Naegleria_gruberi_XP_002674222
Guillardia_theta_CCMP2712_XP_005831207
Cafeteria_roenbergensis_KAA0154571
Porcisia_hertigi_KAG5509145
Trypanosoma_cruzi_KAF5222213
Trypanosoma_cruzi_RNC37465
Angomonas_deanei_EPY32482
Leptomonas_seymouri_KPI85006
Leptomonas_pyrrhocoris_XP_015655420
Triplophysa_rosa_KAI7814089
Danionella_translucida_TRY90509
Clupea_harengus_XP_031440983
Trichoplax_adhaerens_XP_002117363
Mytilus_californianus_XP_052106846
Capsaspora_owczarzaki_ATCC_30864_XP_004363728
Monosiga_brevicollis_MX1_XP_001749400
Phytophthora_kernoviae_RLN50435
Phytophthora_boehmeriae_KAG7398327
Pythium_brassicae_nom_inval_TYZ61647
Nannochloropsis_salina_CCMP1776_TFJ85322
Ectocarpus_sp_CCAP_1310/34_CAB1117158
Benjaminiella_poitrasii_KAI9480974
Cokeromyces_recurvatus_XP_051386473
Mucor_circinelloides_1006PhL_EPB92713
Pilaira_anomala_KAI9354434
Lichtheimia_corymbifera_JMRCFSU9682_CDH49480
Dichotomocladium_elegans_KAI9315508
Hesseltinella_vesiculosa_ORX53550
Diversispora_epigaea_RHZ77343
Thamnocephalis_sphaerospora_RKP05012
Lophiotrema_nucula_KAF2107164
Polyplosphaeria_fusca_KAF2735995

J) pATPase

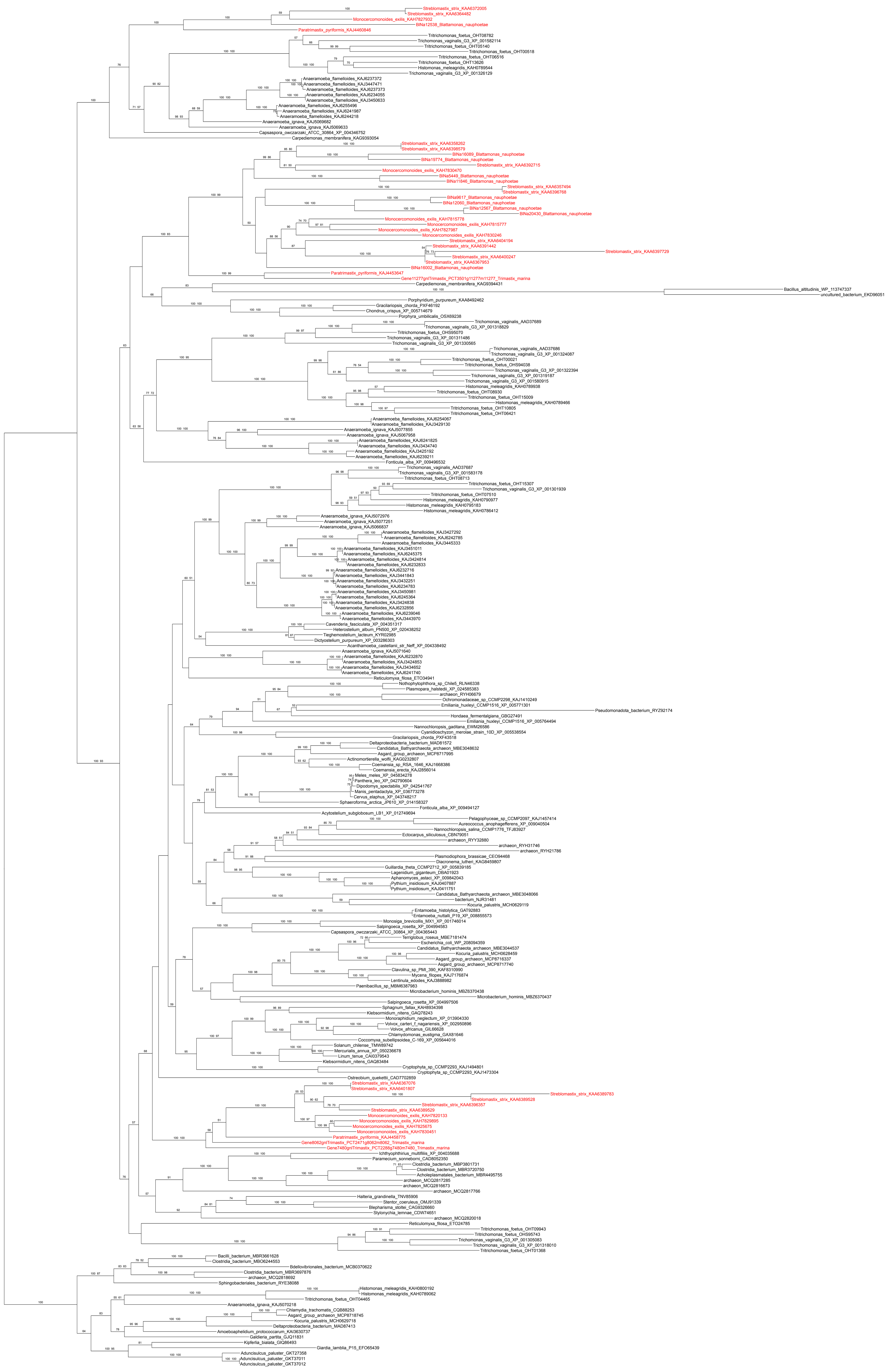

K) SP

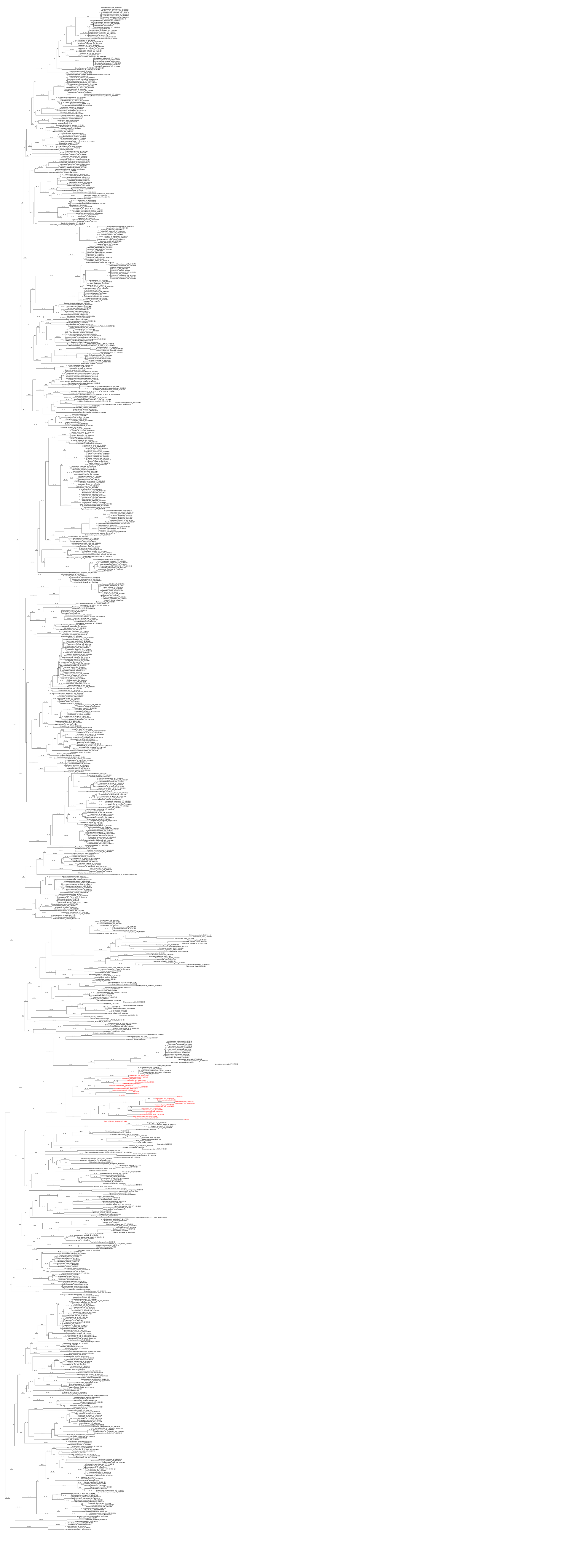
