## Supplementary figures and images for "Genomics of Preaxostyla Flagellates Illuminates the Path Towards the Loss of Mitochondria"

### S1Fig

#### *Paratrimastix pyriformis*

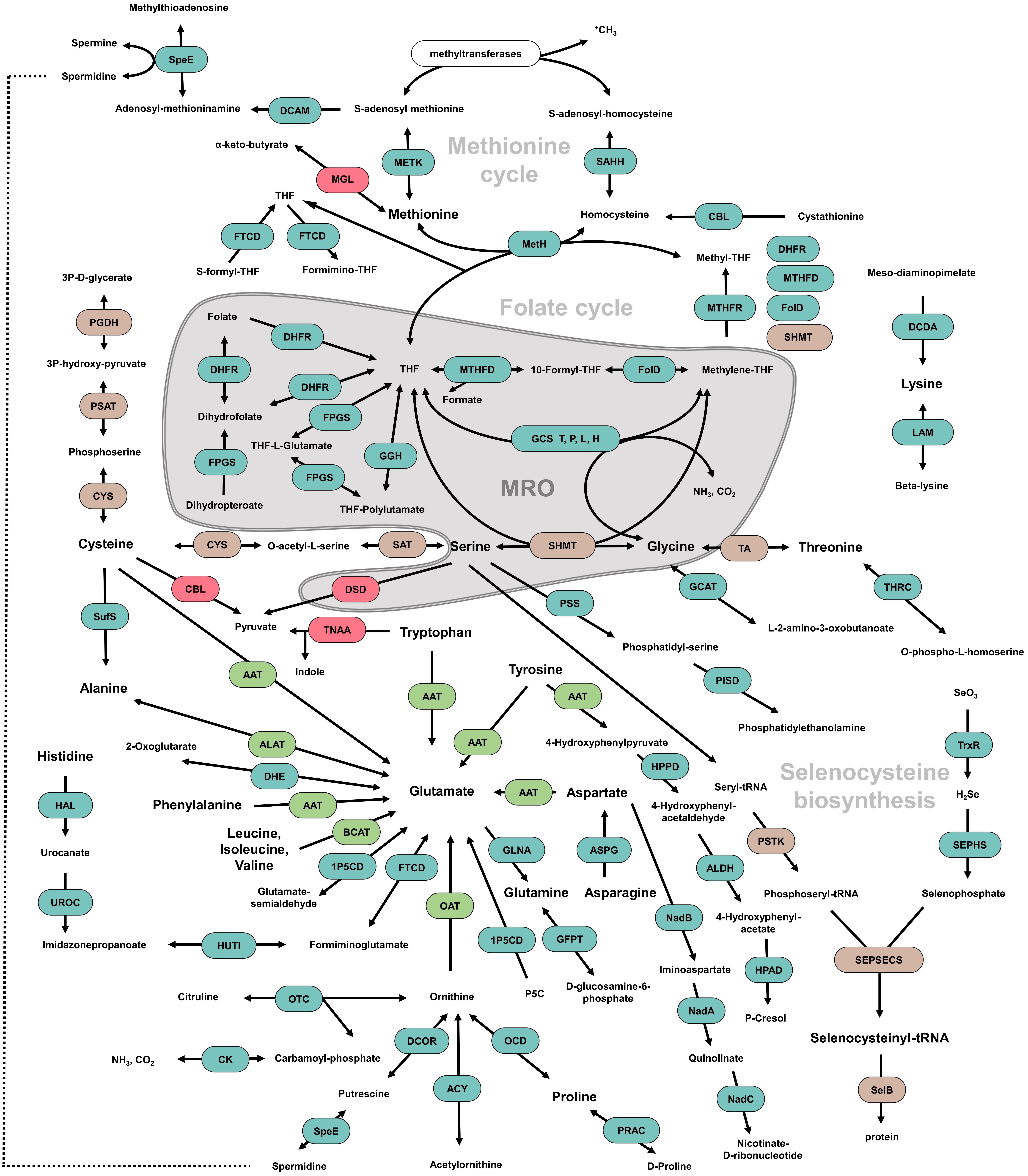

### S2Fig

#### *Trimastix marina*

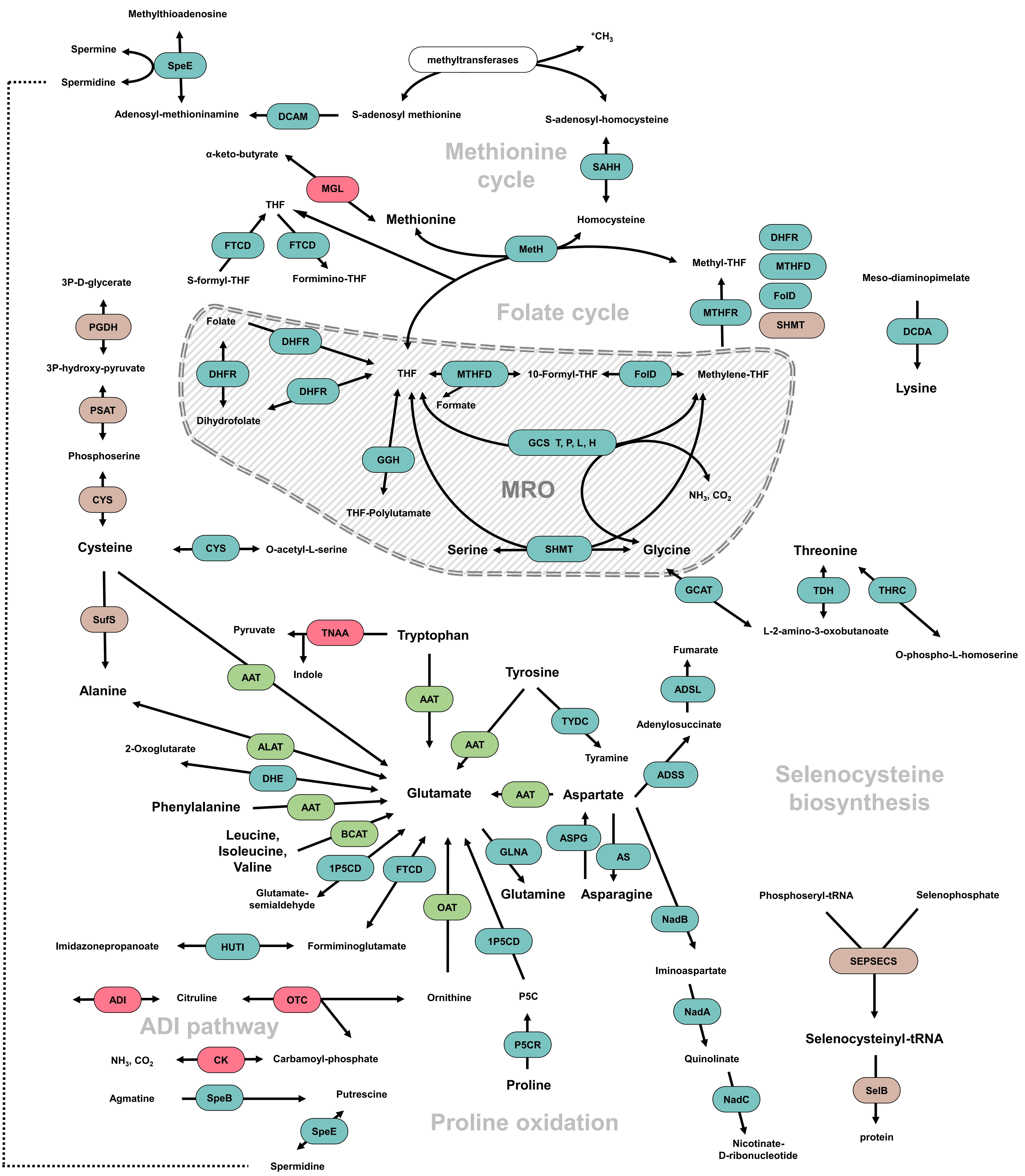

### S3Fig

##### *Monocercomonoides exilis*

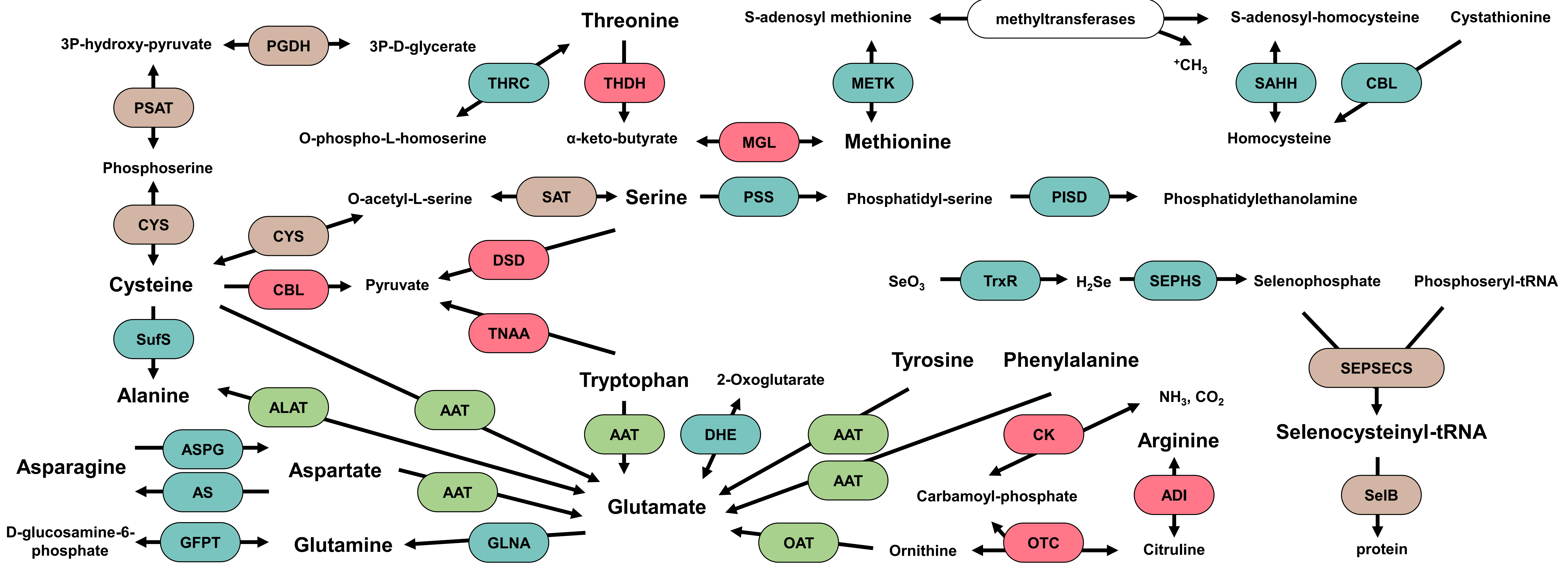


##### *Blattamonas nauphoetae*

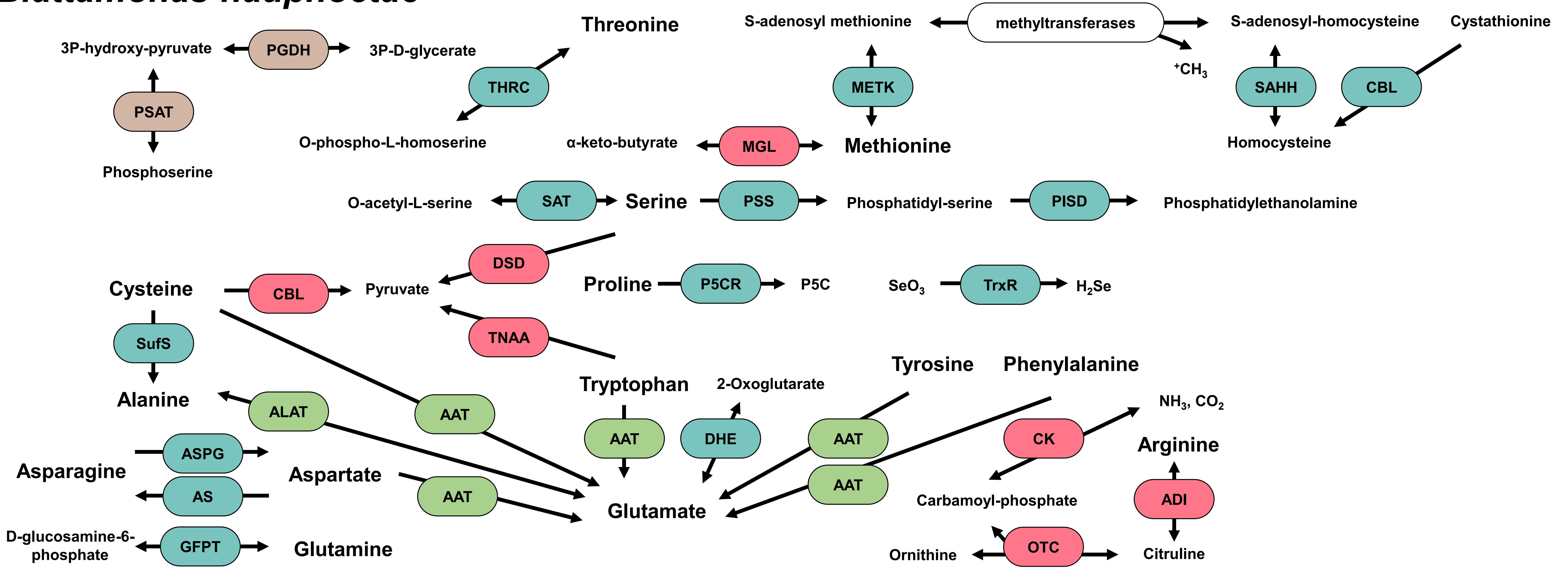


##### *Streblomastix strix*

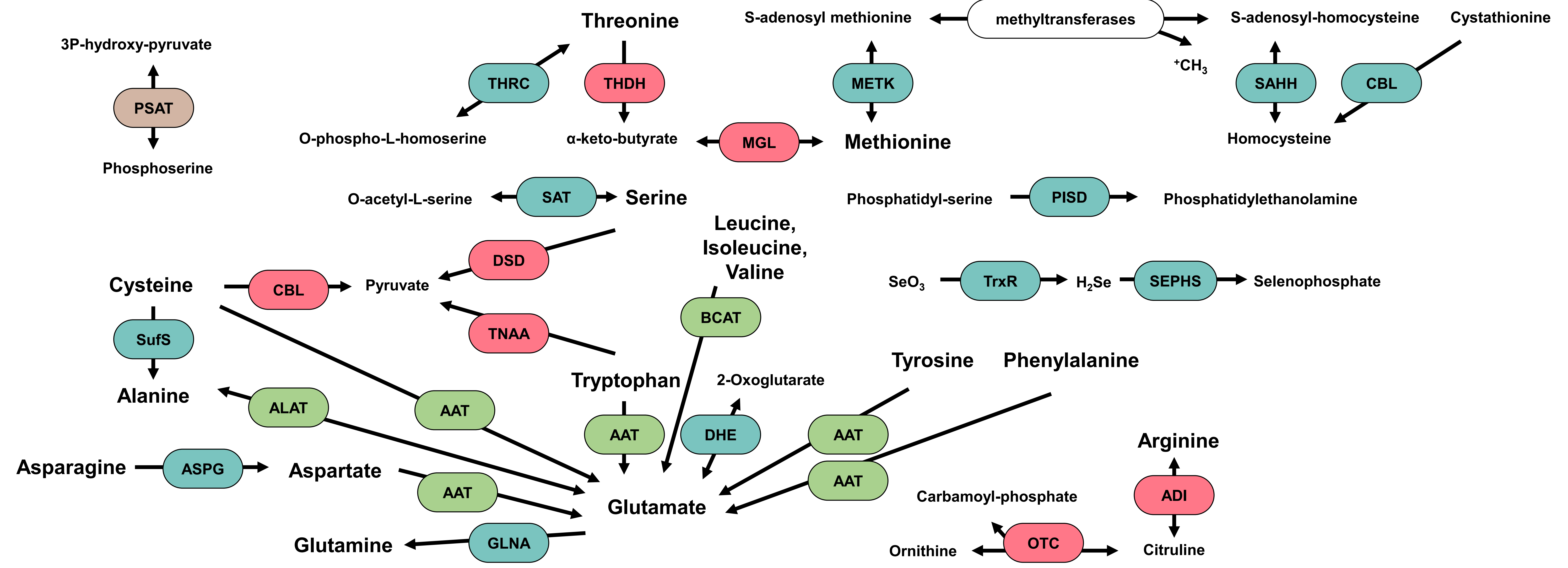

### S4Fig

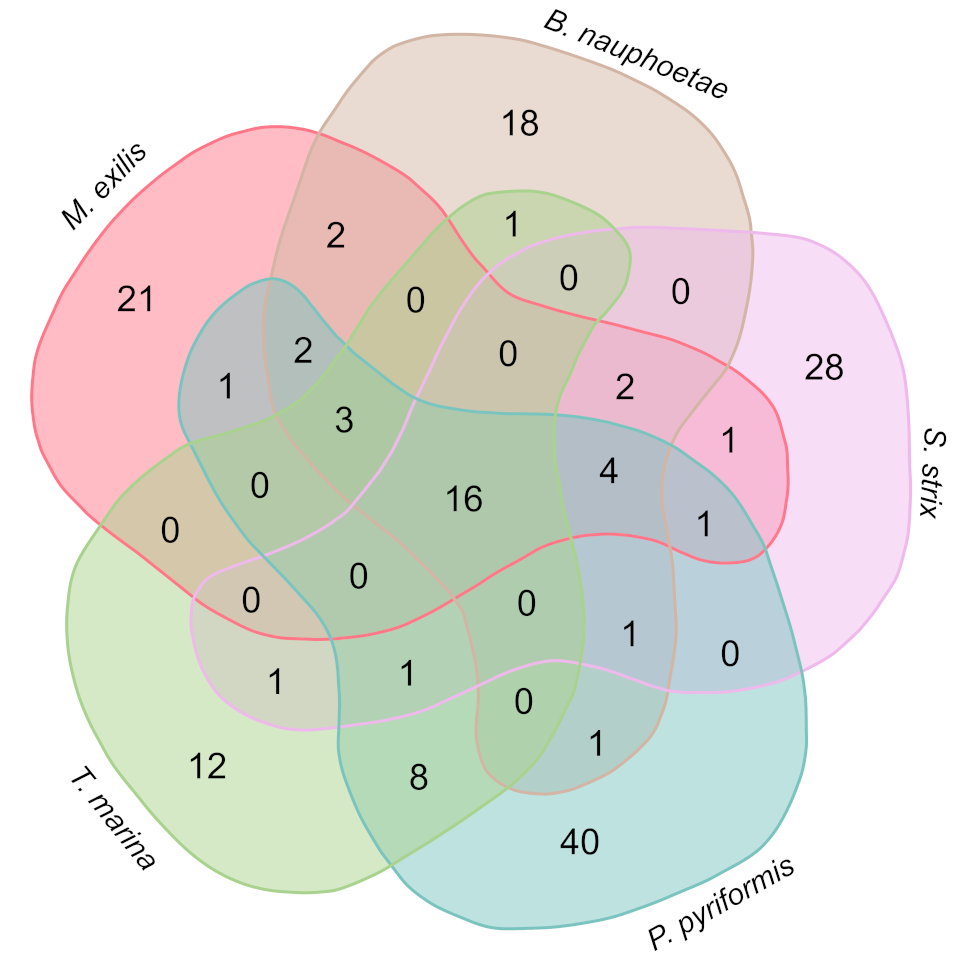
